# RLP23 is required for Arabidopsis immunity against the grey mould pathogen *Botrytis cinerea*

**DOI:** 10.1101/2020.05.03.075408

**Authors:** Erika Ono, Kazuyuki Mise, Yoshitaka Takano

## Abstract

Necrosis- and ethylene-inducing-like proteins (NLPs) are secreted by fungi, oomycetes and bacteria. Conserved nlp peptides derived from NLPs are recognized as pathogen-associated molecular patterns (PAMPs), leading to PAMP-triggered immune responses. RLP23 is the receptor of the nlp peptides in *Arabidopsis thaliana*; however, its actual contribution to plant immunity is unclear. Here, we report that RLP23 is required for Arabidopsis immunity against the necrotrophic fungal pathogen *Botrytis cinerea*. Arabidopsis *rlp23* mutants exhibited enhanced susceptibility to *B. cinerea* compared with the wild-type plants. Notably, microscopic observation of the *B. cinerea* infection behaviour indicated the involvement of RLP23 in pre-invasive resistance to the pathogen. *B. cinerea* carried two *NLP* genes, *BcNEP1* and *BcNEP2*; *BcNEP1* was expressed preferentially before/during invasion into Arabidopsis, whereas *BcNEP2* was expressed at the late phase of infection. Importantly, the nlp peptides derived from both BcNEP1 and BcNEP2 induced the production of reactive oxygen species in an RLP23-dependent manner. In contrast, other necrotrophic fungus *Alternaria brassicicola* did not express the *NLP* gene in the early infection phase and exhibited no enhanced virulence in the *rlp23* mutants. Collectively, these results strongly suggest that RLP23 contributes to Arabidopsis pre-invasive resistance to *B. cinerea* via NLP recognition at the early infection phase.

## Introduction

Plants activate immunity against pathogenic microorganisms through their perception of pathogen-associated molecular patterns (PAMPs), for protection against pathogen infection^1^. Many types of PAMPs have been reported, such as flg22 (derived from bacterial flagellin), elf18 (derived from the bacterial elongation factor-Tu), chitin (derived from fungal cell wall), and the nlp peptides derived from secreted proteins termed necrosis- and ethylene-inducing-like proteins (NLPs), conserved in a broad range of fungi, bacteria and oomycetes^2–6^. PAMPs are recognized by corresponding pattern-recognition receptors (PRRs) localized on the plasma membrane of plant cells^1^. For instance, the flg22 is recognized by a leucine-rich repeat receptor-like kinase (LRR-RK) termed FLAGELLIN SENSITIVE 2 (FLS2)^3^. FLS2 interacts with its co-receptor, BRASSINOSTEROID INSENSITIVE 1-ASSOCIATED KINASE 1 (BAK1), and the two factors trans-phosphorylate each other after perception of flg22^7,8^. A series of phosphorylation events lead to the subsequent activation of PAMP-triggered immune responses, such as reactive oxygen species (ROS) burst, mitogen-activated protein kinase (MAPK) activation and callose deposition^9^. Regarding the limited PRR genes, it was reported that the deletion of a single of these genes reduces the resistance against particular pathogens. For example, Arabidopsis *fls2* mutants are more susceptible to the bacterial pathogen *Pseudomonas syringae* pv. *tomato* DC3000, as assessed via spray-inoculation of the *fls2* mutants with a suspension of the pathogen^10^. The *Arabidopsis* CHITIN ELICITOR RECEPTOR KINASE 1 (CERK1) is the lysin motif receptor-like kinase (LysM-RK) for chitin, and *cerk1* mutants exhibit enhanced susceptibility to the necrotrophic fungal pathogen *Alternaria brassicicola*^5^.

The nlp peptides were identified as a PAMP derived from NLPs^2,6^. The first purified NLP protein, NEP1, was originally isolated from culture filtrates of *Fusarium oxysporum* f. sp. *erythroxyli*, which is the fungus that causes vascular wilt disease in *Erythroxylum coca*^11^. NEP1 induces necrosis and the production of the plant hormone ethylene in plants. It was also reported that NLPs, including NEP1, induce necrosis in eudicot, but not monocot, plants^11–13^. Although many NLPs are cytotoxic to plants, some non-cytotoxic NLPs have been identified in fungi and oomycetes in recent years^14–16^. Subsequent studies revealed that NLPs are categorized into three distinct types, i.e., types I, II and III^17–19^. Type I NLPs are the most widely conserved NLPs. They are present in fungi, bacteria and oomycetes. In contrast, type II NLPs are mainly found in bacteria and fungi, and very few are observed in oomycetes. Type III NLPs are mainly observed only in a limited number of ascomycete species^19^. Subsequent structural analysis of an NLP from the phytopathogenic oomycete *Phytium aphanidermatum* revealed that NLPs have two opposing antiparallel ß-sheets, called ß-sandwich, in the central part of the protein, similar to lectins in fungi and actinoporins in sea anemones^20^. Moreover, it was recently reported that NLPs specifically bind to glycosyl inositol phosphoryl ceramide (GIPC)^21^, as actinoporins bind to sphingomyelin. Although GIPC is the most abundant class of sphingolipids in plants, GIPCs also occur in fungi and protozoa^22^. GIPC is composed of inositol phosphoceramide and is anchored in the membrane^23^. It is speculated that the conformational changes of GIPC–NLP complexes induce pore formation and lead to cell death^21,24^.

Importantly, two independent articles reported that the amino acids that are conserved in the central part of NLPs (a 20-amino-acid pattern termed nlp20 and a 24-amino-acid pattern termed nlp24) act as a PAMP. The nlp24 peptide contains two conserved regions; conserved region I starts with the AIMY sequence of amino acids, which are highly conserved in type I NLPs, whereas conserved region II starts with the heptapeptide motif GHRHDWE, which is highly conserved in all NLPs^6,18^. Treatment with the synthetic nlp20/nlp24 peptide induces defence responses, including ethylene production, in *A. thaliana*^2,6^. The recognition of nlp20 is also observed in other plants of the Brassicaceae family, i.e., *Arabis alpina*, *Thlaspi arvense* and *Draba rigida*, and the Asteraceae family, i.e., *Lactuca sativa*^2^. In contrast, it was reported that cucurbits recognize the C-terminal amino acids of NLPs^25^. Recently, an LRR-receptor protein (LRR-RP) termed RLP23 was identified as the receptor for nlp20 in *A. thaliana*^26^. It was also reported that RLP23 constitutively interacts with the LRR-RK called SUPPRESSOR of BIR1-1 (SOBIR1) and forms a complex with BAK1 after nlp20 perception. The complex triggers the subsequent activation of the nlp peptide-induced immunity^26–28^.

However, the actual contribution of RLP23 to plant immunity remains unclear. Here, we report that RLP23 was required for Arabidopsis immunity against the necrotrophic fungal pathogen *Botrytis cinerea*, which is the causal agent of grey mould. Microscopic observation revealed that the invasion ratio of *B. cinerea* was higher in Arabidopsis *rlp23* mutants compared with wild-type (WT) plants, suggesting the involvement of RLP23 in pre-invasive immunity, which inhibits pathogen invasion. *B. cinerea* carries two *NLP* genes termed *BcNEP1* and *BcNEP2*. In the interaction with Arabidopsis, *BcNEP1* was preferentially expressed before/during pathogen invasion, whereas *BcNEP2* was expressed at the late infection phase. We also discovered that the nlp peptides derived from BcNEP1 and BcNEP2 activated the PAMP-triggered immune response in an RLP23-dependent manner. Together with further studies on *A. brassicicola*, these results strongly suggest that NLP perception via RLP23 in the early infection phase contributes to Arabidopsis immunity against *B. cinerea*.

## Results

### Arabidopsis *rlp23* mutants show enhanced susceptibility to the necrotrophic pathogen *Botrytis cinerea*

Arabidopsis *rlp23* mutants were inoculated with the *Botrytis cinerea* IuRy-1 strain via dropping of a conidial suspension of the pathogen. We found that the Arabidopsis *rlp23-1* plants showed enhanced susceptibility to *B. cinerea* at 7 days post-inoculation (7 dpi) (Fig. 1A); enhanced susceptibility was also observed in Arabidopsis *rlp23-2* plants (Fig. 1A). Lesion size was significantly increased in the *rlp23-1* and *rlp23-2* mutants compared with the parental WT plant (Col-0) (Fig. 1B). A quantitative analysis of the growth of *B. cinerea* in planta also supported this finding. The amount of *B. cinerea* in inoculated Arabidopsis plants was quantified by quantitative PCR (qPCR) of the *B. cinerea* cutinase A gene (*BcCutA*) at 6, 12 and 16 hours post-inoculation (hpi)^29^. The expression of *BcCutA* increased gradually in the inoculated *Arabidopsis* plants from 6 to 16 hpi (Fig. 1C). Importantly, the amount of the amplified *BcCutA* at 12 hpi in the *rlp23-2* mutant was significantly higher than that detected in the WT plants (Fig. 1C). These results suggest that RLP23 contributes to Arabidopsis immunity against the necrotrophic fungal pathogen *B. cinerea*.

**Figure 1.**
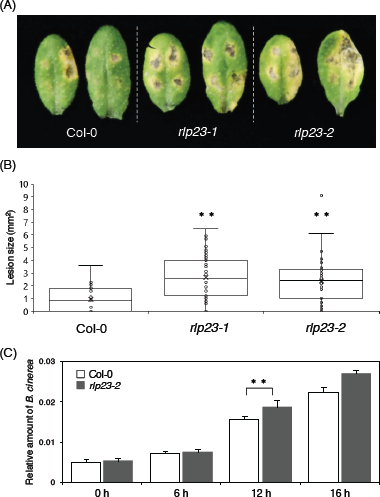
The Arabidopsis *rlp23* mutations enhanced the susceptibility to *B. cinerea*. 4 to 5-week-old plants were inocul ated with 5 µl of conidial suspensions (1 x 10^5^ conidia/mL) of *B. cinerea*. (A) Lesion development on each mutant. The photograph was acquired at 7 dpi. (B) Lesion areas were measured at 7 dpi. At least 24 lesions were measured from each line. The statistical significance of differences in lesion size was determined by Tukey’s honestly significant difference (HSD) test (\*\**P* < 0.01). The experiment was repeat ed three times, with similar results. (C) Growth of *B. cinerea* in planta based on qPCR using *B. cinerea*- and Arabidopsis-specifi c primers. *BcCutA* was quantified by qPCR in the extracted genomic DNA. The Arabidopsis α-shaggy kinase gene (*AtASK)* was used as a reference. Means and SDs were cal culated from three independent samples. The statistical analysis was conducted using two-tailed Student’s *t*-tests. The level of *BcCutA* was compared at the same time points between Col-0 and the *rlp23-2* mutant plants (\*\**P* < 0.01).

### The Arabidopsis *rlp23* mutant has a defect in pre-invasive resistance against *B. cinerea*

To identify the pathogen infection steps that are affected by the loss of RLP23 function, we performed microscopic observation of the *B. cinerea* infection behaviour in Arabidopsis plants. At 6 hpi, most conidia had germinated and started to elongate germ tubes, but did not develop invasive hyphae in WT Col-0 or the *rlp23-2* mutant. At 12 hpi, some of the germinating conidia exhibited invasive hyphae inside plants (Fig. 2A). Specifically, about 25% of germinating conidia in WT plants displayed invasive hyphae at 12 hpi (Fig. 2B). In contrast, about 45% of germinating conidia in the *rlp23-2* mutant showed invasive hyphae at the same time point (Fig. 2B). Thus, the invasion ratio of *B. cinerea* was clearly higher in the *rlp23-2* mutant compared with WT plants at 12 hpi. Considering the higher amount of *B. cinerea* detected at 12 hpi in the *rlp23-2* mutant compared with the wild type (Fig. 1C), these results suggest that the enhanced susceptibility to *B. cinerea* observed in the Arabidopsis *rlp23* mutants is at least partially attributable to the reduction in pre-invasive resistance against the pathogen.

**Figure 2.**
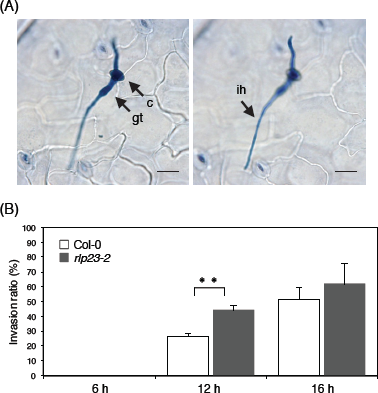
Increased invasion ratio in Arabidopsis *rlp23-2* mutants vs Col-0 plants at 12 hpi. 4 to 5-week-old plants were drop-inoculated with *B. cinerea* (1 x 10^5^ conidia/mL) and collected at each time point. The inoculated leaves were stained with trypan blue solution and the *B. cinerea* hyphae were observed under a light microscope. (A) Arabidopsis Col-0 plants were inoculated with *B. cinerea* and collected at 12 hpi. The image on the left is focused on conidium and the image on the right is focused on invasive hypha. Bars, 20 µm. c: conidium, g: germ tube, ih: invasive hypha. (B) Inoculated leaves were collected at each time point and stained with trypan blue solution. At least 100 appressoria were investigated to determine whether they had developed invasive hyphae. The means and SDs were calculated from three independent samples. The statistical analysis was conducted using two-tailed Student’s *t*-tests. The invasion ratio was compared at the same time points between Col-0 and the *rlp23-2* mutant plants (\*\**P* < 0.01). The experiment was repeated twice, with similar results.

### nlp peptides derived from two types of *B. cinerea* NLPs induce Arabidopsis immunity in an RLP23-dependent manner

Because RLP23 is required for the perception of nlp peptides derived from NLPs and the subsequent activation of immunity^26^, next we investigated whether the reduced immunity against *B. cinerea* observed in the *rlp23* mutants is related to the RLP23-dependent recognition of NLP proteins secreted by *B. cinerea.* It was previously reported that *B. cinerea* secretes two types of NLPs, i.e., BcNEP1 and BcNEP2, during infection of tomato (Fig. 3)^12^. To examine whether *B. cinerea* also secretes these two NLPs during infection of *A. thaliana*, we sprayed a conidial suspension of *B. cinerea* on Arabidopsis plants and quantified the expression level of the *BcNEP1* and *BcNEP2* genes by quantitative reverse transcription PCR (RT–qPCR). *BcNEP2* was preferentially expressed in the late phase of *B. cinerea* infection, which is in accordance with the expression pattern of *NLP* genes in other fungal pathogens^30–32^ (Fig. 4A). The expression of *BcNEP2* was clearly induced at 72 hpi and was highest at 96 hpi. In contrast, the expression level of *BcNEP1* was highest at 12 hpi, a time point at which *B. cinerea* had started to invade, suggesting the preferential expression of *BcNEP1* at the early infection phase (Fig. 4A). Specifically, the expression of *BcNEP1* reached its peak at 12 hpi, decreased to 48 hpi, and then increased again from 48 hpi to 96 hpi (Fig. 4A). This result seems to be consistent with that reported previously for tomato^12^. Thus, in Arabidopsis, although *B. cinerea* expresses *BcNEP2* at the late infection phase, the pathogen expresses *BcNEP1* at high levels before (6 hpi) and during (12 hpi) invasion, which is distinct from the case of *BcNEP2*. These findings suggest that the perception of BcNEP1 plays a key role in the pre-invasive resistance of *A. thaliana* against *B. cinerea*. We also investigated the expression of *RLP23* during *B. cinerea* infection and found that *RLP23* was constitutively expressed, although it started to be induced to some degree at 24 hpi (Fig. 4B).

**Figure 3.**
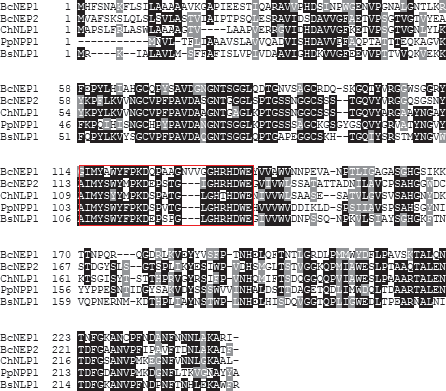
BcNEP1, BcNEP2, ChNLP1, PpNPP1 and BsNLP1 exhibited a high similarity in amino acid sequence. Sequence data of *B. cinerea* NEP1 (BcNEP1) and NEP2 (BcNEP2), *C. higginsianum* NLP1 (ChNLP1), *P. parasitica* NPP1 (PpNPP1) and *B. subtilis* NLP1 (BsNLP1) can be found in the GenBank/EMBL data libraries under the accession numbers XP_001555180 for BcNEP1, XP_001551049 for BcNEP2, XP_018154754 for ChNLP1, AAK19753.1 for PpNPP1 and WP_019714591.1 for BsNLP1. Amino acid sequences were aligned using Clustal/Omega^33^. The putative nlp24 region is indicated by the red open box.

**Figure 4.**
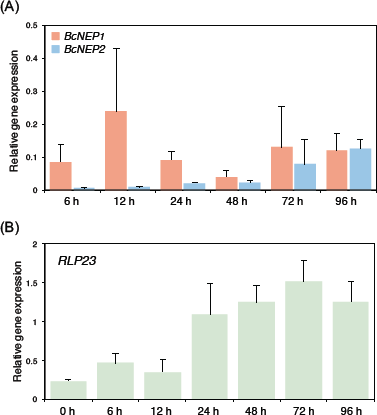
*BcNEP1* was preferentially expressed in the early phase of *B. cinerea* infection. A conidial suspension (3 x 10^5^ conidia/mL) of *B. cinerea* was spray-inoculated onto Col-0 plants and total RNA was extracted. (A) The transcripts of *BcNEP1* and *BcNEP2* were quantified by RT–qPCR. *BcUBQ* was used as a housekeeping gene. Means and SDs were calculated from three independent samples. The experiment was repeated twice, with similar results. (B) *RLP23* was quantified by RT– qPCR. *AtUBC* was used as a housekeeping gene. Means and SDs were calculated from three independent samples. The experiment was repeated twice, with similar results.

We then wondered whether BcNEP1 was actually recognized by Arabidopsis RLP23. Focusing on the amino acid sequences of BcNEP1 and BcNEP2, we performed multiple alignment of the amino acid sequences of BcNEP1, BcNEP2, and ChNLP1 of *Colletotrichum higginsianum*, PpNPP1 of *Phytophthora parasitica* and BsNLP1 of *Bacillus subtilis* using Clustal/Omega^2,18,33^. This analysis showed that these proteins had a high similarity in their amino acid sequences (Fig. 3). However, in the case of the predicted nlp24 region, BcNEP1 exhibited a difference vs. the other four NLPs of microorganisms in three kingdoms. Although the nlp24 region of the five NLPs, including BcNEP1, commonly possessed two conserved regions, i.e., conserved region I and conserved region II, the corresponding sequence of BcNEP1 contained additional three amino acids (NVV) between conserved region I and conserved region II (Fig. 5A).

**Figure 5.**
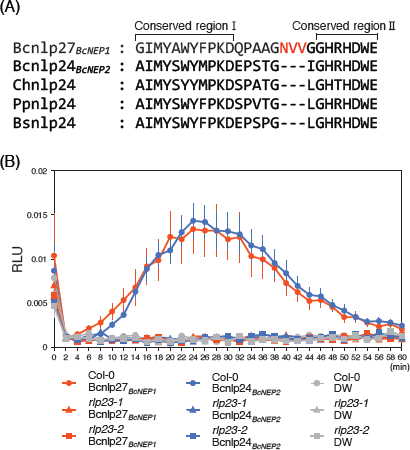
Bcnlp27 derived from BcNEP1 and Bcnlp24 derived from BcNEP2 triggered ROS accumulation in Arabidopsis in an RLP23-dependent manner. (A) nlp27 from BcNEP1 was aligned with nlp24 from BcNEP2, ChNLP1, PpNPP1 and BsNLP1. The three additional amino acids located between the conserved region I and conserved region II in Bcnlp27 are highlighted by red characters. (B) Leaf discs from true leaves of 4–5-week-old Arabidopsis Col-0, *rlp23-1* and *rlp23-2* mutant plants were treat ed with 500 nM Bcnlp27_*BcNEP1*_, 500 nM Bcnlp24_*BcNEP2*_ or sterile distilled water (DW). Data are reported as relative light units (RLU) and represent the mean ± SE (n = 8). The experiment was repeated three times, with similar results.

The 20 and 24 amino acids derived from the nlp sequence of BcNEP2 were reported to activate Arabidopsis immunity^2,6^; however, it remains unknown whether the corresponding 27 amino acids derived from BcNEP1 (termed Bcnlp27_*BcNEP1*_ here) can activate immunity. Here, we named the 24 amino acids derived from the BcNEP2 nlp sequence as Bcnlp24_*BcNEP2*_. Because Bcnlp27_*BcNEP1*_ contains three additional amino acids in the nlp24 sequence, we were not able to exclude the possibility that the insertion blocks its recognition by Arabidopsis RLP23. To test whether Bcnlp27_*BcNEP1*_ can be recognized by RLP23 and trigger immunity, we measured the amount of ROS after treatment of Arabidopsis leaf discs with each synthetic nlp peptide. We found that Bcnlp24_*BcNEP2*_ induced ROS production, as reported previously (Fig. 5B). Notably, Bcnlp27_*BcNEP1*_ also induced ROS production, to the same extent as did Bcnlp24_*BcNEP2*_ (Fig. 5B). In contrast, ROS production was suppressed in the *rlp23* mutants (Fig. 5B). These results demonstrate that not only Bcnlp24_*BcNEP2*_ but also Bcnlp27_*BcNEP1*_ triggers the PAMP-triggered immune response in *A. thaliana* in an RLP23-dependent manner, supporting the idea that the recognition of BcNEPs, especially BcNEP1, by RLP23 at the early infection phase contributes to Arabidopsis pre-invasive immunity against *B. cinerea*.

### The necrotrophic pathogen *Alternaria brassicicola* does not express the *NLP* gene at the early infection phase and does not exhibit enhanced virulence in *rlp23* plants

*A. brassicicola* is a necrotrophic fungal pathogen that induces lesions in Arabidopsis leaves, similar to *B. cinerea*^34^. Therefore, next we investigated whether the *A. brassicicola* strain Ryo-1 also expresses its *NLP* gene at the early infection phase in Arabidopsis. Via the alignment of the *A. brassicicola* genome sequence with the *A. alternata* mRNA sequence using Exonerate^35^, we found that *A. brassicicola* carries only one *NLP* homologue in its genome sequence (Supplementary Fig. 1). We designated this gene *AbNLP1* and quantified its expression by RT–qPCR during *A. brassicicola* infection. We observed that *AbNLP1* was expressed at very low levels at 24 hpi, increased to 48 hpi and then decreased towards 72 hpi (Fig. 6A). Importantly, *AbNLP1* expression was not detected at 4 and 12 hpi. We also found that the *A. brassicicola* strain Ryo-1 starts to invade Arabidopsis at around 12 hpi^36^. Collectively, these results indicate that *AbNLP1* of *A. brassicicola* is not expressed before/during invasion, in contrast to *BcNEP1* of *B. cinerea.* Next, to determine whether the lack of *RLP23* also reduces the immunity of Arabidopsis against *A. brassicicola*, a conidial suspension of the *A. brassicicola* strain was drop-inoculated on the *rlp23* mutants. The *rlp23-1* and *rlp23-2* mutants exhibited immunity against *A. brassicicola* at 4 dpi, similar to that observed in the WT plants (Fig. 6B) and in contrast to the case of *B. cinerea* inoculation. The size of the lesions caused by *A. brassicicola* in the *rlp23-1* and *rlp23-2* mutants was not significantly different from that of Col-0 (Fig. 6C), whereas the *cyp71A12 cyp71A13* mutant exhibited enhanced susceptibility to the pathogen, consistent with our recent finding^36^. Thus, RLP23 is not essential for the Arabidopsis immunity against *A. brassicicola*.

**Figure 6.**
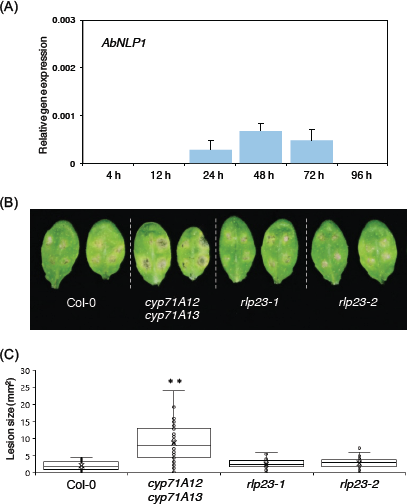
The Arabidopsis *rlp23* mutation did not enhance the susceptibility to *A. brassicicola*. (A) A conidial suspension (5 x 10^5^ conidia/mL) of *A. brassicicola* was spray-inoculated onto 4 to 5-week-old Col-0 plants. *AbNLP1* was quantified by RT–qPCR. *AbEF1* was used as a housekeeping gene. Means and SDs were calculated from three independent samples. *AbNLP1* was only detected at 24, 48 and 72 hpi in Arabidopsis Col-0 plants. (B) Leaves from 4–5-week-old plants were drop-inoculated with 5 µl of conidial suspensions (1 x 10^5^ conidia/mL) of *A. brassicicola.* The susceptibility to *A. brassicicola* was not affected in Arabidopsis *rlp23-1* and *rlp23-2* mutants compared with Col-0 plants, whereas the *cyp71A12 cyp71A13* mutant exhibited enhanced susceptibility to this pathogen. The photograph was taken at 4 dpi. (C) Lesion areas were measured in the experiments (B). At least 40 lesions from each line were measured at 4 dpi. The statistical significance of differences in lesion size was determined by Tukey’s honestly significant difference (HSD) test (\*\**P* < 0.01). The experiment was repeated twice, with similar results.

We also investigated the possible role of RLP23 toward a fungal pathogen taking infection strategies that are distinct from those of *B. cinerea*. *C. higginsianum* is a hemibiotrophic fungal pathogen, i.e., the pathogen initially establishes a biotrophic infection in host cells, which is followed by a necrotrophic phase that leads to cell death and the emergence of pathogenic lesions^37^. The host plant species of *C. higginsianum* is *Brassica rapa*, and it also infects *A. thaliana*^38^. We inoculated *C. higginsianum* (MAFF305635) on the Arabidopsis *rlp23* plants by dropping a conidial suspension of the pathogen and measured the lesion size at 5 dpi. The *rlp23-1* and *rlp23-2* mutants did not exhibit enhanced susceptibility to *C. higginsianum*, and there was no significant difference compared with Col-0 regarding the size of the lesions caused by *C. higginsianum* (Supplementary Fig. 2). These results showed that RLP23 is of particular importance to the immunity against *B. cinerea*, whereas it is dispensable for the immunity against *A. brassicicola* and *C. higginsianum*.

### Bcnlp peptides enhance the Arabidopsis immunity against *A. brassicicola*

The results described above strongly suggest that the perception of Bcnlp27_*BcNEP1*_ of BcNEP1 at the early infection phase contributes to Arabidopsis immunity against *B. cinerea*. To address this contention further, we inoculated *A. brassicicola* on Arabidopsis WT plants by dropping a conidial suspension together with the Bcnlp27_*BcNEP1*_ peptide. We found that Arabidopsis plants that were inoculated with the pathogen and Bcnlp27_*BcNEP1*_ concomitantly showed enhanced resistance against *A. brassicicola* compared with plants that were inoculated with the pathogen alone (Fig. 7A). The lesion size caused by *A. brassicicola* was significantly reduced in the Bcnlp27_*BcNEP1*_-treated plants compared with the plants that were inoculated with *A. brassicicola* alone (Fig. 7B). Similar results were obtained for the co-inoculation of *A. brassicicola* and the Bcnlp24_*BcNEP2*_ peptide (Fig. 7). These results demonstrate that the activation of the nlp-induced immunity in the early infection phase strengthens the Arabidopsis immunity against *A. brassicicola*.

**Figure 7.**
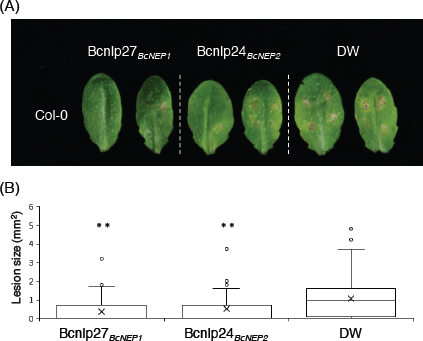
Inoculation of *A. brassicicola* together with the Bcnlp peptides reduced its virulence in Arabidopsis Col-0 plants. Leaves from 4–5-week-old plants were drop-inocul ated with 5 µl of conidial suspensions (1 x 10^5^ conidia/mL) of *A. brassicicola* together with 500 nM Bcnlp27_*BcNEP1*_, 500 nM Bcnlp24_*BcNEP2*_ or sterile distilled water (DW). (A) Lesion development caused by *A. brassicicola*. The photograph was taken at 5 dpi. (B) Lesion areas were measured in the experiments (A). At least 60 lesions were measured from each line at 5 dpi. The statistical significance of differences in lesion size was determined by Tukey’s honestly significant difference (HSD) test (\*\**P* < 0.01). The experiment was repeated twice, with similar results.

## Discussion

*RLP23* encodes an RLP that is essential for the perception of the nlp24 peptides and subsequent activation of immune responses. However, the actual contribution of RLP23 to the Arabidopsis immunity against pathogens remains poorly understood. In this study, we revealed that RLP23 was involved in Arabidopsis immunity against the necrotrophic fungal pathogen *B. cinerea* (Fig. 1). In contrast, RLP23 was dispensable for the immunity against the necrotrophic fungus *A. brassicicola* and the hemi-biotrophic fungus *C. higginsianum* (Figs. 6B, 6C and Supplementary Fig. 2). Interestingly, microscopic observation revealed that the invasion ratio of *B. cinerea* was increased in the *rlp23* mutant compared with WT plants (Fig. 2B). Furthermore, *BcNEP1* was expressed before/during invasion, in contrast to that observed for *BcNEP2*, which was expressed at the late infection phase (Fig. 4A). Furthermore, not only the nlp sequence of BcNEP2 (Bcnlp24_*BcNEP2*_) but also that of BcNEP1 (Bcnlp27_*BcNEP1*_) was recognized by RLP23, although the Bcnlp27_*BcNEP1*_ sequence contained three additional amino acids that were absent in BcNEP2 and in the NLPs of *C. higginsianum*, *P. parasitica* (oomycete) and *B. subtilis* (bacterium) (Fig. 5A). Furthermore, we found that *A. brassicicola* expressed its *NLP* gene (*AbNLP1*) preferentially at the late infection phase, rather than before/during pathogen invasion (Fig. 6A). We also showed that the co-inoculation of *A. brassicicola* with the Bcnlp27_*BcNEP1*_ peptide strengthened the immunity against *A. brassicicola* (Fig. 7). Collectively, these results revealed that RLP23 contributes to the Arabidopsis preinvasive resistance to *B. cinerea* via the recognition of the NLP proteins, mainly BcNEP1, which are secreted at the early infection phase.

In hemi-biotrophic pathogens, including Colletotrichum fungi, it was proposed that NLP proteins function in the transition from the biotrophic phase to the necrotrophic phase and in the subsequent maintenance of the necrotrophic phase^32^. This idea is consistent with the findings that *NLP* expression is restricted to the late infection phase of hemi-biotrophic pathogens^31,39,40^. However, our previous work on *C. orbiculare*–cucurbit interactions suggested another explanation for the restriction of *NLP* expression to the late phase^25^. We generated a transgenic strain of *C. orbiculare* that expressed the *NLP* gene not only at the late infection phase but also at the early infection phase, and found that the generated *C. orbiculare* transformants failed to infect cucurbits by activating their pre-invasive resistance^25^. In this case, cucurbits did not recognize the nlp24 region; rather, they recognized the C-terminal region of the NLP protein^25^.

The results reported here provide the first example of the activation of pre-invasive resistance by a WT fungal pathogen, rather than an artificial transgenic pathogen^25^, via plant recognition of the pathogen NLP protein, which further supports the relationship between the restriction of NLP expression to the late phase and the avoidance of pathogen NLP recognition by plants at the early phase. Our study also provides evidence that supports the idea that PRR-dependent PAMP recognition contributes to pre-invasive resistance to fungal pathogens in higher plants. This finding is also consistent with the knowledge that non-host plant resistance against a broad range of fungal pathogens largely relies on pre-invasive immune responses^41–45^.

Because Arabidopsis recognizes BcNEP1 via RLP23, the expression of *BcNEP1* at the early infection phase has a negative impact on *B. cinerea* infection. Why does *B. cinerea* express *BcNEP1* at the early infection phase? Does *B. cinerea* benefit from *BcNEP1* expression? We consider that *B. cinerea* expresses *BcNEP1* to induce cell death in host plants at the early infection phase, thus facilitating its infection, because *B. cinerea* is a necrotrophic fungal pathogen. Interestingly, BcNEP1 reportedly induces necrosis more efficiently compared with BcNEP2^13^. However, a single-gene knockout mutant of *BcNEP1* was previously generated and the diameters of the lesions caused by the Δ*Bcnep1* mutant were not significantly different from those caused by the *B. cinerea* WT strain in detached tomato and *Nicotiana benthamiana* leaves^12^. Thus, there is currently no direct evidence that *BcNEP1* actually contributes to the *B. cinerea* virulence. Also, it is noteworthy that the nlp24 peptide is recognized by a limited number of plant species, including several Brassicae species^2^. Therefore, we speculate that *B. cinerea* evolved to express *BcNEP1* at the early phase to cause cell death in host plants that are not able to recognize BcNEP1, because *B. cinerea* fungi generally exhibit a broad host range.

The RLP23-triggered antifungal immune pathways that are crucial for pre-invasive resistance against *B. cinerea* also remain to be elucidated. A recent report revealed that the nlp24 peptide strongly induces ethylene production in an RLP23-dependent manner compared with flg22^26^. Ethylene is a plant hormone that regulates diverse developmental and physiological processes, including fruit ripening, seed germination, abscission, senescence and immunity to pathogens^46^. In particular, ethylene contributes to immunity against necrotrophic pathogens, including *B. cinerea*, compared with other types of pathogens^47^. For example, ethylene insensitive 2 (EIN2) is an endoplasmic reticulum (ER) membrane-localized positive regulator of ethylene, and the Arabidopsis *ein2* mutants are more susceptible to *B. cinerea* infection^48,49^. Constitutive expression of *ERF1*, an early ethylene responsive gene^50^, also enhances the Arabidopsis resistance against *B. cinerea*^51^. Therefore, we speculate that the induction of ethylene production by BcNEP1 might be involved in the activation of antifungal immune responses that restrict the entry of *B. cinerea*. Further studies are necessary to elucidate at the molecular level the preinvasive immune responses that are triggered by the RLP23-dependent nlp recognition.

In contrast to that observed for *B. cinerea*, the *rlp23* mutants showed no enhanced susceptibility to *A. brassiciacola* (Figs. 6B and 6C). It is likely that RLP23 did not contribute to the immunity against *A. brassiciacola* because of (i) the late timing of *AbNLP1* expression and (ii) the low expression level of *AbNLP1* (too low to activate immunity). Interestingly, the *ein2* mutants did not exhibit enhanced susceptibility to *A. brassicicola*, unlike that observed for *B. cinerea*, even though *A. brassicicola* is also a necrotrophic pathogen^52^. Alternatively, the Arabidopsis immune responses that are effective against *A. brassicicola* might be distinct from those that are effective against *B. cinerea*. However, we showed that the co-inoculation of *A. brassicicola* with the Bcnlp peptides enhanced the Arabidopsis immunity to this pathogen (Fig. 7). Therefore, we assume that the immune responses triggered by the nlp peptides are commonly effective against *B. cinerea* and *A. brassicicola.*

## Materials and Methods

### Plant growth

Seeds of *A. thaliana* were sown on soil or rockwool (Grodan), incubated for 2 days at 4°C in the dark and then grown at 22°C in 16 h daylight with water for soil or Hoagland medium for rockwool.

### *Arabidopsis* T-DNA insertion lines

All mutant lines used in this study were derived from Col-0. The *rlp23-1*^26^ and *rlp23-2*^26^ mutants were provided by the Arabidopsis Biological Resource Center (ABRC), Ohio University, USA, and the *cyp71A12 cyp71A13* mutant^53^ was provided by Dr. Paweł Bednarek, Polish Academy of Sciences, Poland.

### Fungal materials

The *B. cinerea* strain IuRy-I was provided by Dr. Katsumi Akutsu, Ibaraki University, Japan. The *A. brassicicola* strain Ryo-1 was provided by Dr. Akira Tohyama. The *C. higginsianum* isolate MAFF305635 was obtained from the Ministry of Agriculture, Forestry and Fisheries (MAFF) Genebank, Japan. *B. cinerea* was cultured on 3.9% (w/v) potato dextrose agar (PDA, Difco) medium at 24°C in the dark. For sporulation of *B. cinerea*, the fungus was cultured under a cycle of 16 h black light and 8 h dark for 4–5 days. *A. brassicicola* and *C. higginsianum* were cultured on 3.9% PDA medium (Nissui) at 24°C in the dark.

### Pathogen inoculation, lesion development analysis and trypan blue staining assay

4 to 5-week-old plants grown on rockwool were used for the inoculation assay. Conidial suspensions (5 µl) of *B. cinerea* (1 x 10^5^ conidia/mL), *A. brassicicola* (1 x 10^5^ conidia/mL) or *C. higginsianum* (1 x 10^5^ conidia/mL) were placed onto each leaf. In the case of *B. cinerea* inoculation, conidia were dissolved in Sabouraud maltose broth buffer [1% (w/v) peptone (Difco) and 4% maltose were dissolved in distilled water and the pH was adjusted to 5.6 with HCl]^54^. The inoculated plants were kept at high humidity and transferred to a growth chamber with 21°C light and 18°C dark temperatures with a 12 h light/12 h dark cycle. For *A. brassicicola* and *C. higginsianum*, conidia were dissolved in sterile distilled water and the inoculated plants were kept at high humidity and at 22°C in 16 h daylight.

For the analysis of lesion development after the inoculation assay, four drops of 5 µl of a conidial suspension of each pathogen were placed onto each leaf, and at least 24 lesions were evaluated in each experiment. The developed lesions were quantified using the ImageJ image analysis software (http://imagej.net). For pathogen invasion assay, trypan blue staining of inoculated leaves was conducted according to Koch and Slusarenko (1990)^55^. For the trypan blue assay, at least 200 conidia were investigated in each treatment area.

### Quantitative RT–PCR analysis and quantitative PCR analysis

For the quantification of the amount of *B. cinerea*, 4–5-week-old plants grown on rockwool were drop-inoculated with *B. cinerea* (1 x 10^5^ conidia/mL) and three leaves were collected. Total DNA was extracted using a DNeasy Plant Mini Kit (Qiagen). For the quantification of gene expression levels during pathogen infection, 4–5-week-old plants grown on rockwool were spray-inoculated with *B. cinerea* (3 x 10^5^ conidia/mL) or *A. brassicicola* (5 x 10^5^ conidia/mL) and three leaves were collected. Total RNA was extracted using an RNeasy Plant Mini Kit (Qiagen). The Takara Prime Script^TM^ RT Master Mix (Takara Bio Inc.) was used for cDNA synthesis.

Takara TB Green^TM^ Premix Ex Taq^TM^ I and a Thermal Cycler Dice Real Time System TP800 (Takara) were used for both quantitative RT–PCR and quantitative PCR using the primers listed in Supplementary Table 1. For quantitative RT–PCR, the *Botrytis* ubiquitin (*BcUBQ*, XM_001556819.1), *Arabidopsis* ubiquitin-conjugating enzyme 2 (*AtUBC*, At5g25760) and *A. brassicicola* elongation factor 1 (*AbEF1*, GEMY01015044) genes were used as internal controls to normalize the levels of cDNA. For quantitative PCR, *Arabidopsis* α-shaggy kinase (*AtASK*, At5g26751) was used as the control. The primers used for the amplification of *BcCutA*, *AtASK*^29^, *AtUBC*^56^, *BcUBQ*^57^ and *AbEF1*^58^ were as described previously.

### Synthetic peptides

The synthetic peptides used in this study were as follows: Bcnlp27_*BcNEP1*_, GIMYAWYFPKDQPAAGNVVGGHRHDWE; and Bcnlp24_*BcNEP2*_, AIMYSWYMPKDEPSTGIGHRHDWE. Each peptide was dissolved in sterile distilled water.

### ROS measurements

The ROS assay was performed as described previously, with some modifications^59^. Leaf discs from true leaves of 4–5-week-old plants grown on soil were kept in the dark overnight with 50 µl of sterile distilled water in a 96-well plate. A reaction solution (50 μl) including 400 µM luminol, 20 µg/ml of horseradish peroxidase and 1 µM synthetic peptides (Bcnlp27_*BcNEP1*_ and Bcnlp24_*BcNEP2*_) was added into each well just before measurement. Luminescence was measured for about 60 min every 2 min using Luminoskan Ascent (Thermo Fisher Scientific). At least eight leaf discs were measured per experimental plot.

## Acknowledgements

We thank Dr. Paweł Bednarek (Polish Academy of Sciences, Poland) for *cyp71A12 cyp71A13,* Arabidopsis Biological Resource Center (Ohio University, USA) for *rlp23-1*, and *rlp23-2*, Dr. Katsumi Akutsu (Ibaraki University, Japan) for *B*. *cinerea* IuRy-1, Dr. Akira Tohyama for *A. brassicicola* Ryo-1 and Ministry of Agriculture, Forestry and Fisheries Genebank (Japan) for *C. higginsianum* MAFF305635. This work was supported by Grants-in-Aid for Scientific Research (18H02204, 18H04780, 18K19212) (KAKENHI), by grants from the Project of the NARO Bio-oriented Technology Research Advancement Institution (Research program on development of innovative technology), and by the Asahi Glass Foundation.

## Authors Contributions

Y.T. and E.O. designed this research. E.O performed the experiments and analyzed the data. E.O., Y.T. and K.M. wrote the manuscript and prepared the figures.

## Notes

### Competing Interest Statement

The authors have declared no competing interest.

