## Supplemental Information for "RLP23 is required for Arabidopsis immunity against the grey mould pathogen *Botrytis cinerea*"

**Supplementary Table 1.** The list of primers used for qPCR

| Gene Name | | Sequence |
|---|---|---|
| *BcCutA* | Forward | AGCCTTATGTCCCTTCCCTTG |
| | Reverse | GAAGAGAAATGGAAAATGGTGAG |
| *AtASK* | Forward | CTTATCGGATTTCTCTATGTTTGGC |
| | Reverse | GAGCTCCTGTTTATTTAACTTGTACATACC |
| *BcNEP1* | Forward | CCATACAGTGCCGTTGATGG |
| | Reverse | GTTTGGCCCTTGCTCTGATC |
| *BcNEP2* | Forward | AAGTCGTAAATGGATGCGTACC |
| | Reverse | CGCTTTGTCCTCCTCGAAC |
| *BcUBQ* | Forward | CAAGGTTACCGACAACAATA |
| | Reverse | GCATCCATCAACTTCTTCAA |
| *AtRLP23* | Forward | GGAGTGGCTTGTCAAGATAATTGG |
| | Reverse | CCCAATTTTATCCTCATTTGCCCG |
| *AtUBC* | Forward | CTGCGACTCAGGGAATCTTCTAA |
| | Reverse | TTGTGCCATTGAATTGAACCC |
| *AbNLP1* | Forward | TGACACTGGGCAAGTTCTCG |
| | Reverse | TGGATTTGCACATCCCTGAG |
| *AbEF1* | Forward | GGGTCCTCGACAAGTTGAA |
| | Reverse | GGGAGCGTCAATAACTGTGA |

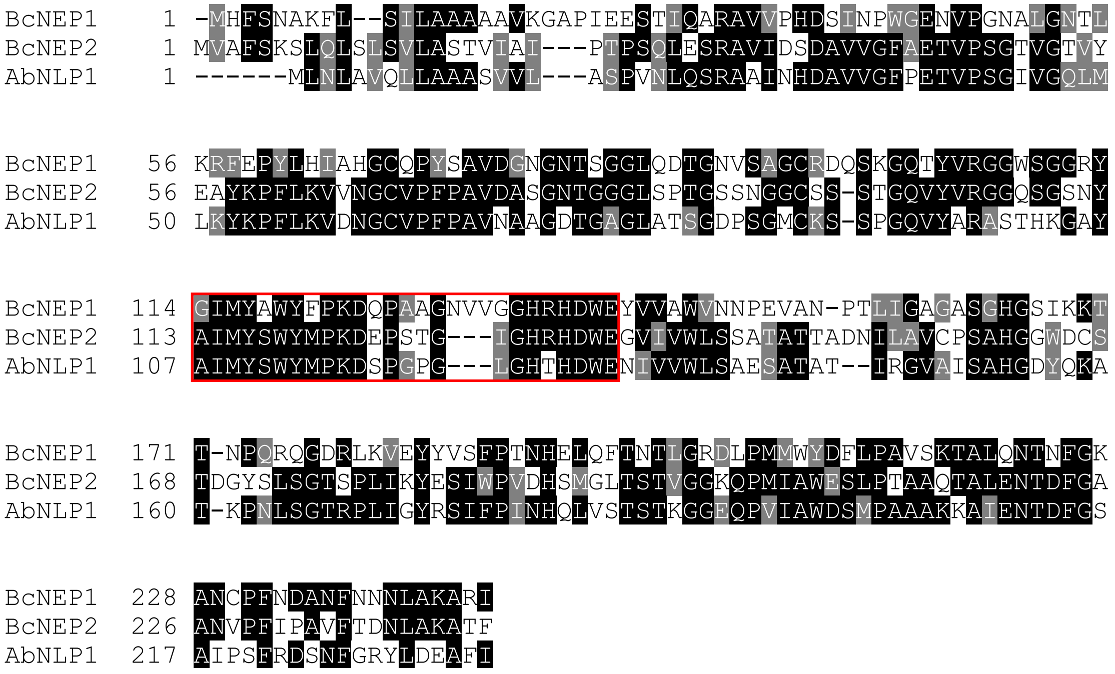


**Supplementary Figure 1. BcNEP1, BcNEP2 and AbNLP1 exhibited a high similarity in amino acid sequence.**

The putative NLP1 homologue in *A. brassicicola* (AbNLP1) was searched in the *A. brassicicola* genome sequence (PHFN01000002.1) by aligning it with the *A. Alternata* mRNA sequence (NW_017306222.1) using Exonerate[35]. Amino acid sequences were aligned using Clustal/Omega[33]. The putative nlp24 region is indicated by the red open box.

(A)

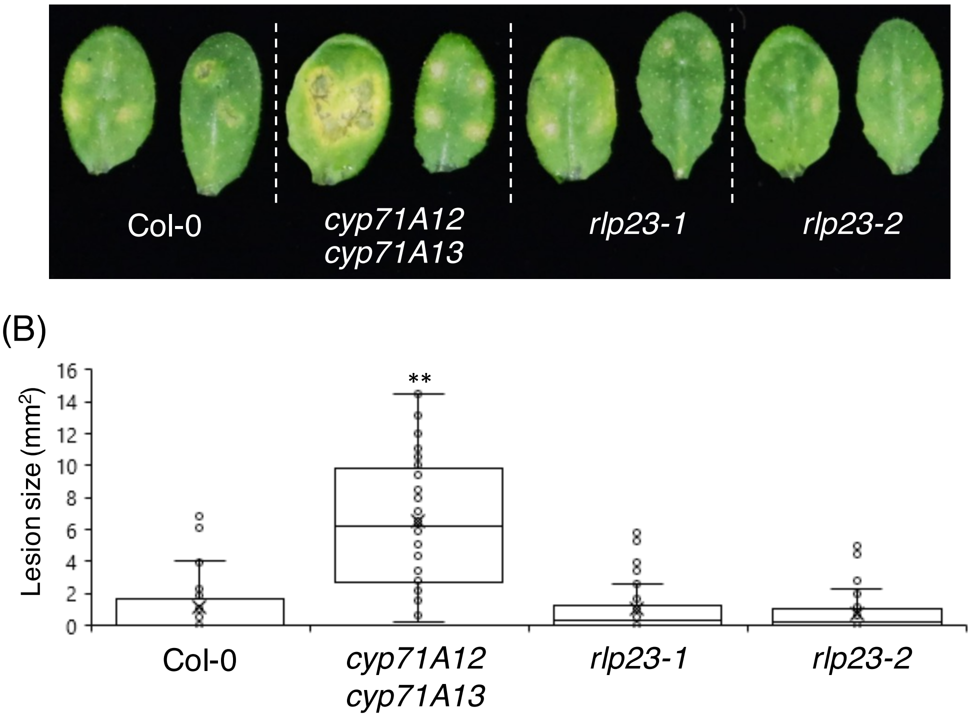


**Supplementary Figure 2. The Arabidopsis *rlp23* mutation did not enhance the susceptibility to *C. higginsianum*.**

4 to 5-week-old plant were inoculated with 5 µl drops of conidial suspension (1 x $10^5$ conidia/mL) of *C. higginsianum*. (A) Lesion development on each mutant caused by *C. higginsianum*. The susceptibility to *C. higginsianum* was not affected in Arabidopsis *rlp23-1* and *rlp23-2* mutants compared with Col-0 plants, whereas the *cyp71A12 cyp71A13* mutant exhibited enhanced susceptibility to the pathogen. The photograph was taken at 5 dpi. (B) Lesion areas were measured in the experiments (A). At least 40 lesions from each line were measured at 5 dpi. The statistical significance of differences in lesion size was determined by Tukey's honestly significant difference (HSD) test (**$P < 0.01$). The experiment was repeated twice, with similar results.
